# Maximum likelihood estimation of the geometric niche preemption model

**DOI:** 10.1101/2021.01.27.428381

**Authors:** Jan Graffelman

**Affiliations:** Department of Statistics and Operations Research, Technical University of Catalonia; Department of Biostatistics, University of Washington

**Keywords:** geometric series, preemption *t* test, broken stick model, rank-abundance plot, robustness

## Abstract

The geometric series or niche preemption model is an elementary ecological model in biodiversity studies. The preemption parameter of this model is usually estimated by regression or iteratively by using May’s equation. This article proposes a maximum likelihood estimator for the niche preemption model, assuming a known number of species and multinomial sampling. A simulation study shows that the maximum likelihood estimator outperforms the classical estimators in this context in terms of bias and precision. We obtain the distribution of the maximum likelihood estimator and use it to obtain confidence intervals for the preemption parameter and to develop a preemption *t* test that can address the hypothesis of equal geometric decay in two samples. We illustrate the use of the new estimator with some empirical data sets taken from the literature and provide software for its use.

## 1 Introduction

The statistical modeling of the relative abundance of a set of species in an ecological community has a longstanding history (Wilson, 1991). Classical elementary models are MacArthur’s (1957) broken stick model, Fisher’s (1943) log series, the geometric model (Motomura, 1932) and the log-normal model (Preston, 1962). Magurran (2004) provides an excellent introduction to these models. Over the last decades, many more refined models have been proposed (Tokeshi, 1990, 1996). Notwithstanding, the most elementary models such as the broken stick model and the geometric series form important references and are widely applied (Fattorini, 2005); they are usually the ones that are first fitted before more complicated alternatives are considered. In this article we focus on the geometric series, also known as the *niche preemption hypothesis*. This model assumes each species succesively exploits a fraction *k* of the available resources, such that the first species exploits fraction *k* of the total resources, the second species fraction *k* of the remaining 1 – *k* resources, and so on. The exploited fraction is reflected by the relative abundance of the species in the community. Mathematically, the model is described by

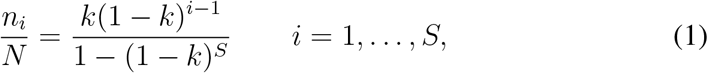

where *N* is the total number of individuals found, *S* the total number of species, *n_i_* the abundance of the *i*th species, and *k* the niche preemption parameter. In our notation, we will use *n*_(*i*)_ to refer to the ordered abundances, such that *n*_(1)_ and *n*_(*S*)_ represent the most and least abundant species respectively. Doi and Mori (2013) gives more historical background on the geometric series. Though the geometric model is regarded as deterministic (Magurran, 2004), we note that the right hand side of Eq. (1) corresponds to the probability function of a truncated geometric distribution. This model implies that the logarithm of the relative abundance decays linearly with the rank of the species, as illustrated in a logarithmic rank-abundance plot in Figure 1 for various values of *k*.

**Figure 1:**
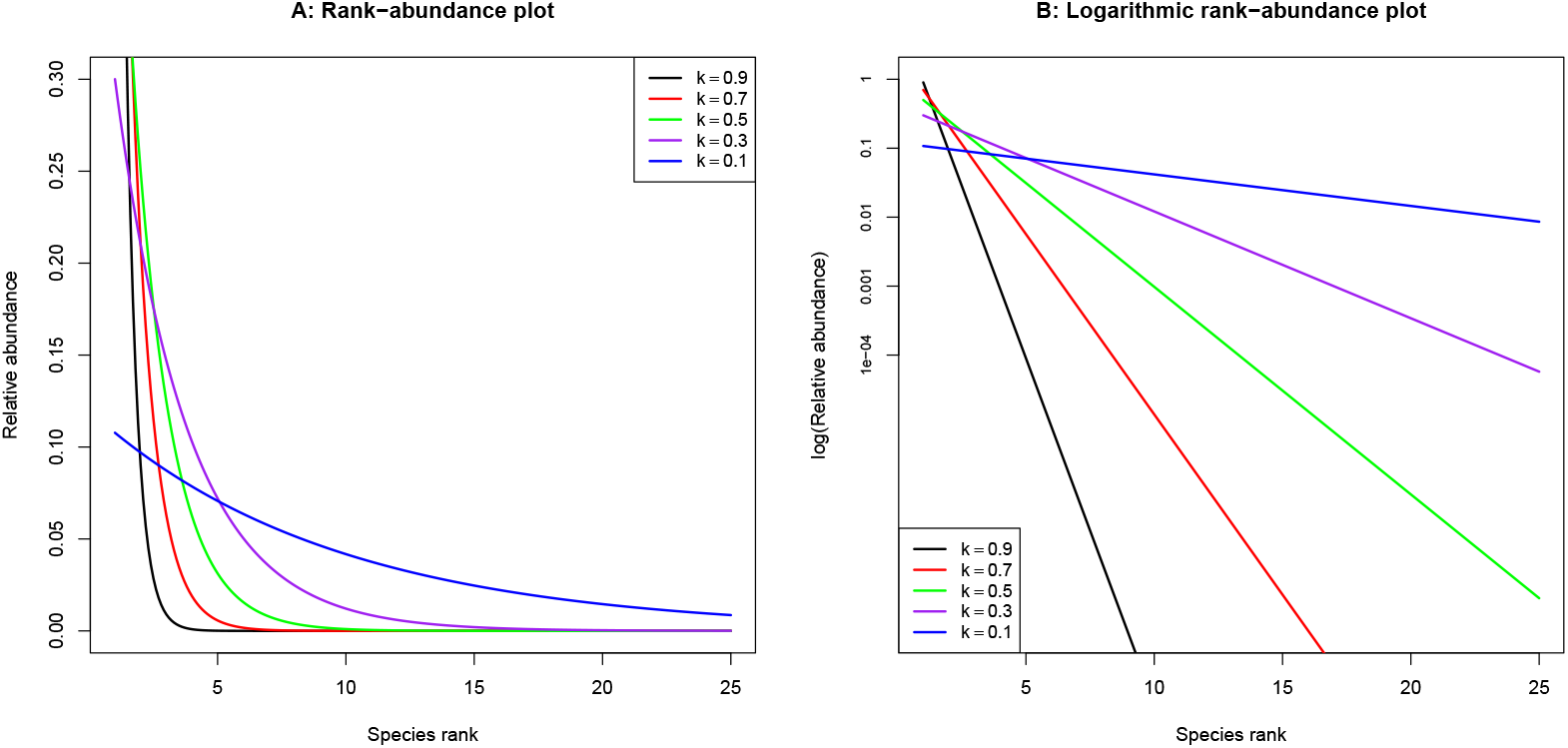
Rank-abundance plots for varying *k* of the geometric model with *S* = 25, with relative scale (panel A) and logarithmic scale (panel B).

The geometric model has been found adequate for species-poor assemblages, resource-poor environments (Fattorini, 2005) and has also been advocated for ecosystems that suffer from anthropogenic disturbance (Caruso and Migliorini, 2006) or that exhibit strong dominance of a few species (Keeley and Fotheringham, 2003). Several estimators for *k* have been proposed in the literature. He and Tang (2008) used several estimators and observed that they provide similar estimates of *k*. However, a statistical study that compares the different estimators assessing their precision and bias seems not available. The main goal of this article is to present a new estimator based on maximum likelihood (ML), and to compare the different estimators in terms of bias and precision. The structure of the article is the following. In Section 2 we review existing estimators for the niche preemption parameter, develop the ML estimator and present the preemption *t* test. In Section 3 we compare the different estimators in a simulation study. Section 4 shows some applications using ecological datasets taken from the literature. We finish with a discussion of the different estimators.

## 2 Preemption parameter estimation and the preemption *t* test

We briefly summarise some popular estimators for *k*, and develop the maximum likelihood estimator. May (1975) proposed to estimate *k* by solving the equation

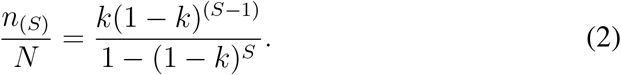

This result is obtained by applying Eq. (1) to the mininum abundance with rank *i* = *S*. This equation can easily be solved on a computer by applying an algorithm searching for the root of a non-linear equation. He and Tang (2008) used both the minimum and maximum abundance, and proposed the estimator

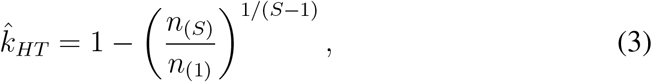

which follows from the fact that successive abundances have a constant ratio. The least-squares regression estimator (He and Tang, 2008), 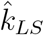, is obtained by noting that the logarithm of the relative abundance is linear in the rank *i* of the species with slope *b*_1_ = ln (1 – *k*);

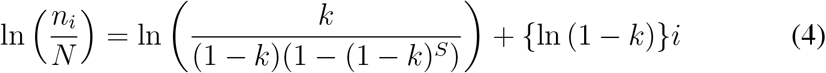

By transforming the slope, we have:

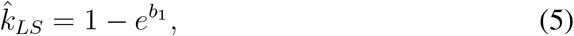

where *b*_1_ is the least squares estimator for the slope obtained by simple linear regression of 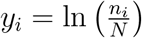 on rank *i*. We note here that 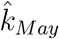 and 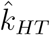 do not assume a statistical model for the data, but employ the geometric series in a completely deterministic manner. The geometric series is considered to be a deterministic niche apportionment model (Magurran, 2004, p. 47). Consequently, there is no estimation of a measure of uncertainty for these estimators. The regression estimator 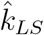 assumes that deviations from geometric decay follow a normal distribution and therefore an expression of the uncertainty of the estimate can be obtained by back-transforming the limits of the confidence interval for *β*_1_:

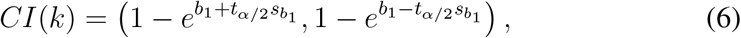

where *s*_*b*_1__ is the standard error of the slope, and *t*_α/2_ the upper percentile of a Student *t* distribution with *S* – 2 degrees of freedom. We proceed by developing the maximum likelihood estimator. If the number of species *S* is considered fixed, then the data consists of counts in a limited number of *S* categories, which can be probabilisitically modeled by the multinomial distribution, given by.

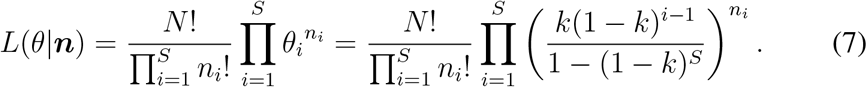

Under the geometric model, the parameters of this multinomial distribution are restricted, such that the parameter *θ_i_* is given by the truncated geometric distribution given by the right hand side of Eq. (1). Maximizing the log-likelihood analytically, we find no closed form solution for *k*. The ML estimator 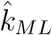 must be found by numerically solving the equation

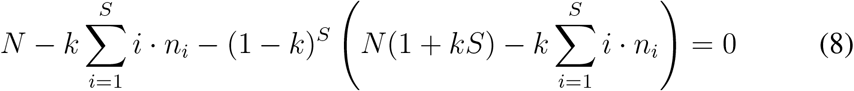

for *k*. For communities with few dominant species (large *k*), the last term in Eq. (8) will generally be small, and the ML estimator can be approximated, in closed form, by

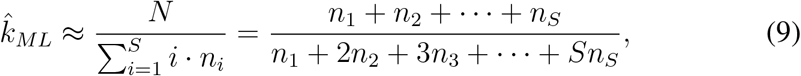

which is the total abundance divided by the abundance-weighted sum of the ranks. Eq. (9) can be used as initial estimate for the fast resolution of Eq. (8). By developing the second derivative of the log-likelihood function, the variance of the ML estimator is obtained as

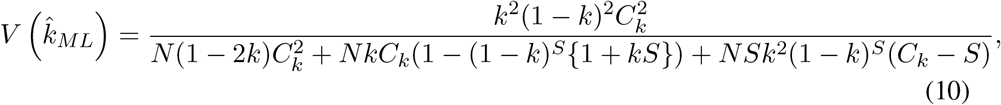

with *C_k_* = 1 – (1 – *k*)^S^, and this variance can be estimated by substituting 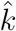 for *k*. Because the ML estimator is asymptotically unbiased, efficient, and normally distributed (Casella and Berger, 2002, Chapter 10), we can construct a 100(1 – *α*) percent confidence interval for *k* which is given by

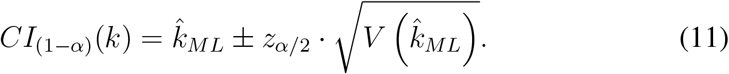

The limits of this confidence interval allow for hypothesis testing with *k*, and they can also be used to show the uncertainty in the estimate of *k* in a rank-abundance plot (See Figures 3 and 4) by painting a corresponding grey area around the line of decay. We study the statistical properties of the new ML estimator and its classical counterparts in a simulation study in Section 3.

Many biodiversity studies are of comparative nature. It is often of interest to compare two (or more) independent samples with respect to some measure of diversity. Such tests have been developed for the Shannon index (Hutcheson, 1970) and for Simpson’s index (Brower et al., 1998), but are apparently not available for the preemption parameter of the geometric series. To test the null hypothesis *k*_1_ = *k*_2_ against *k*_1_ ≠ *k*_2_, we can use the test statistic

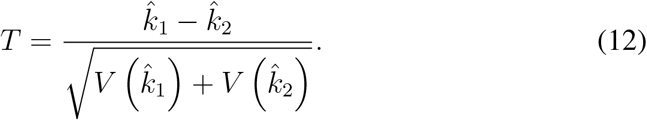

The development of this test is analogous to a standard two-sample *t* test for equality of means without assuming equality of variances for the two groups (DeGroot, 1986), using the Welch modification. Under the null, statistic *T* follows a student *t* distribution with degrees of freedom (df) given by:

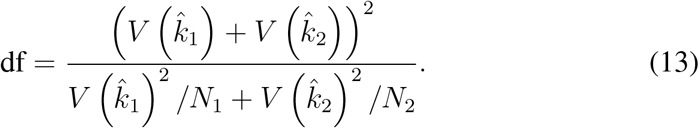

We refer to this test as the *preemption t test*. Some examples are given in Section 4 below. In practice, *N*_1_ and *N*_2_ are large, and the standard normal distribution can be used for the calculation of the p-value.

## 3 Monte Carlo simulations

We simulate species counts by drawing samples from the multinomial distribution given by Eq. (7), for given *N, S* and a considering a sequence of values (0.1, 0.2,… 0.9) for preemption parameter *k*. We repeat simulations 10,000 times, computing all four estimators presented in the previous section. Boxplots of the values of the estimators obtained in the simulations are shown in Figure 2. This figure shows the ML estimator has the smallest variance for all values of *k*. All other estimators typically have more bias than the ML estimator. Table 1 summarizes the results of the simulation, quantifying bias, variance and mean squared error (MSE) for all estimators and different values of *k*.

**Figure 2:**
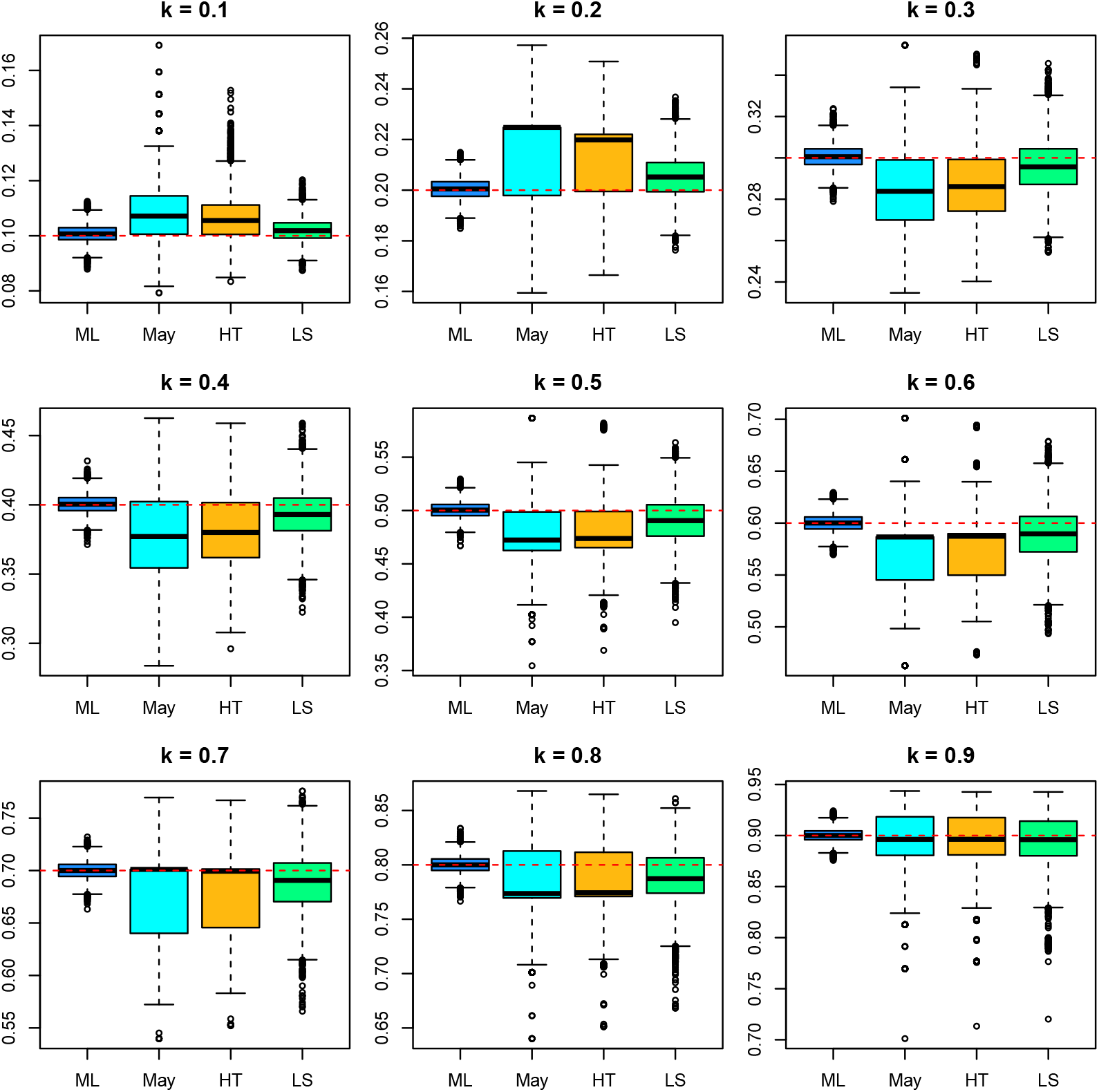
Monte Carlo simulations for the geometric series. Distribution of the different estimators for *N* = 2,000, *S* = 25 and various values of *k*.

**Table 1:** Mean, variance, bias and mean squared error for different estimators of preemption parameter *k*.

| $k$ | Method | mean | bias | var | mse |
| --- | --- | --- | --- | --- | --- |
| 0.1 | ML | 0.1007 | 0.0007 | 0.00001 | 0.00001 |
|  | May | 0.1084 | 0.0084 | 0.00012 | 0.00019 |
|  | HT | 0.1063 | 0.0063 | 0.00007 | 0.00011 |
|  | LS | 0.1020 | 0.0020 | 0.00002 | 0.00002 |
| 0.2 | ML | 0.2005 | 0.0005 | 0.00002 | 0.00002 |
|  | May | 0.2156 | 0.0156 | 0.00025 | 0.00049 |
|  | HT | 0.2132 | 0.0132 | 0.00018 | 0.00035 |
|  | LS | 0.2053 | 0.0053 | 0.00007 | 0.00010 |
| 0.3 | ML | 0.3006 | 0.0006 | 0.00003 | 0.00003 |
|  | May | 0.2862 | -0.0138 | 0.00035 | 0.00054 |
|  | HT | 0.2879 | -0.0121 | 0.00027 | 0.00041 |
|  | LS | 0.2959 | -0.0041 | 0.00016 | 0.00018 |
| 0.4 | ML | 0.4005 | 0.0005 | 0.00005 | 0.00005 |
|  | May | 0.3813 | -0.0187 | 0.00068 | 0.00103 |
|  | HT | 0.3834 | -0.0166 | 0.00053 | 0.00081 |
|  | LS | 0.3930 | -0.0070 | 0.00031 | 0.00035 |
| 0.5 | ML | 0.5005 | 0.0005 | 0.00006 | 0.00006 |
|  | May | 0.4792 | -0.0208 | 0.00102 | 0.00145 |
|  | HT | 0.4814 | -0.0186 | 0.00082 | 0.00117 |
|  | LS | 0.4907 | -0.0093 | 0.00047 | 0.00056 |
| 0.6 | ML | 0.6001 | 0.0001 | 0.00007 | 0.00007 |
|  | May | 0.5788 | -0.0212 | 0.00127 | 0.00172 |
|  | HT | 0.5808 | -0.0192 | 0.00106 | 0.00143 |
|  | LS | 0.5894 | -0.0106 | 0.00063 | 0.00074 |
| 0.7 | ML | 0.7002 | 0.0002 | 0.00007 | 0.00007 |
|  | May | 0.6798 | -0.0202 | 0.00137 | 0.00178 |
|  | HT | 0.6815 | -0.0185 | 0.00117 | 0.00151 |
|  | LS | 0.6891 | -0.0109 | 0.00072 | 0.00084 |
| 0.8 | ML | 0.8001 | 0.0001 | 0.00006 | 0.00006 |
|  | May | 0.7829 | -0.0171 | 0.00129 | 0.00158 |
|  | HT | 0.7841 | -0.0159 | 0.00114 | 0.00139 |
|  | LS | 0.7895 | -0.0105 | 0.00075 | 0.00086 |
| 0.9 | ML | 0.9001 | 0.0001 | 0.00004 | 0.00004 |
|  | May | 0.8896 | -0.0104 | 0.00072 | 0.00083 |
|  | HT | 0.8902 | -0.0098 | 0.00066 | 0.00076 |
|  | LS | 0.8924 | -0.0076 | 0.00048 | 0.00054 |

For small value of *k* (≤ 0.20) the estimators of May, He and Tang, and the regression estimator have positive bias, and for larger values they have negative bias. Table 1 shows that the ML estimator has the smallest bias, variance and MSE in all settings, and is clearly the estimator with the best statistical properties. The least-squares estimator has generally less bias than the estimators of May and He and Tang. The estimator of May has the largest variance, and also presents more outliers. Similar results were obtained for larger and smaller values of *N* (results not shown).

### 4 Analysis of empirical data sets

In this section we apply the different estimators to some empirical datasets taken from the ecological literature. Many data sets are available at the Ecological Register (Alroy, 2015). We use Australian bird abundances (Fattorini, 2005; Magurran, 1988) and Indian dung beetles (Ganeshaiah et al., 1997; Magurran, 2004) to illustrate the difference between estimators of the preemption parameter. We use the dung beetle data from Mehrabi et al. (2014) to illustrate the preemption *t*-test.

#### 4.1 Australian bird abundances

The abundances of *S* = 31 bird species in wet sclerophyll forest, totalling *N* = 834 individuals were recorded. Figure 3 shows the rank-abuance plot of this data, with a fitted line for each of the four estimators discussed in Section 2. The numerical estimates of the preemption parameter are very similar for all four estimators (See Table 2) and by visual inspection the geometric model is seen to fit the data very well. The grey zone in the plot is determined by the confidence limits for the ML estimator. All other estimators give values inside this confidence interval, and can be considered not to differ significantly from the ML estimate.

**Figure 3:**
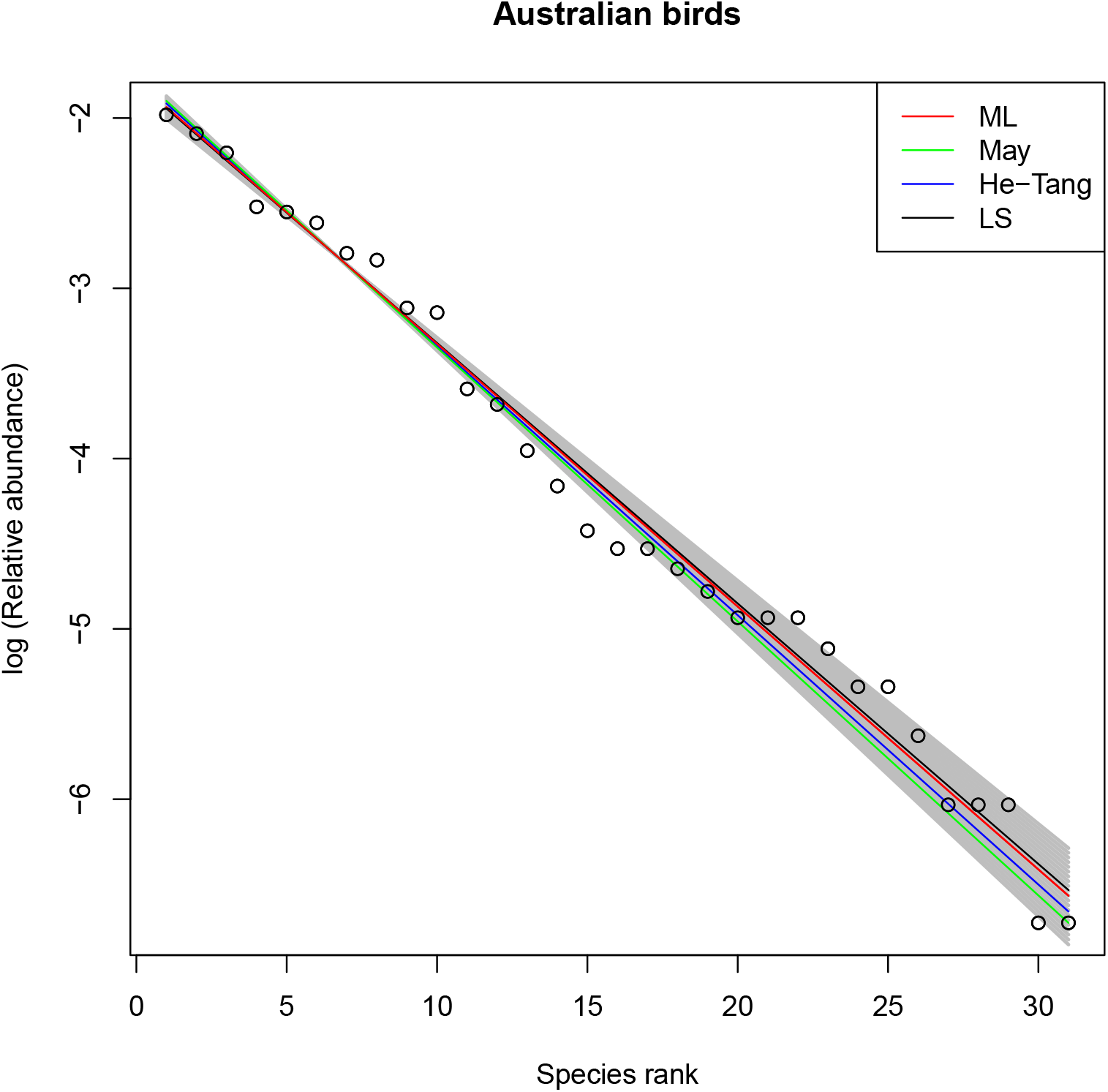
Rank-abundance plot of Australian birds in wet sclerophyll forest. Fitted lines represent geometric models estimated by maximum likelihood (ML), May’s equation (May), He-Tang’s estimator and least-squares regression (LS).

**Table 2:** Estimates of the preemption parameter for the Australian bird data according to different methods (se = standard error, CI = confidence interval).

| Estimator | $\hat{k}$ | se | 95% CI |
| --- | --- | --- | --- |
| May | 0.149 | - | - |
| HT | 0.146 | - | - |
| LS | 0.142 | - | (0.136, 0.148) |
| ML | 0.143 | 0.0051 | (0.133, 0.153) |

#### 4.2 Indian dung beetles

Figure 4 shows the rank-abundance plot of the Indian dung beetle data. Note that there is a considerable difference between May’s classical estimator and the ML estimate. The ML estimator is 31% larger. Expected relative frequencies (in the log scale) have been calculated and plotted in Figure 4 to show the fit of all estimators. This shows May’s estimator underestimates the frequency of the most abundant beetle, and overestimates the frequencies of almost all other species. The ML estimator fits the abundant species much better and is seen to underestimate the rare species. The values of the different estimators are given in Table 3. We note that May’s classical estimator, He-Tang’s estimator and the regression estimator are all outside the confidence interval of the ML estimator. There is clearly a significant difference between the ML estimator and its alternatives, the ML estimator suggesting a stronger decay.

**Figure 4:**
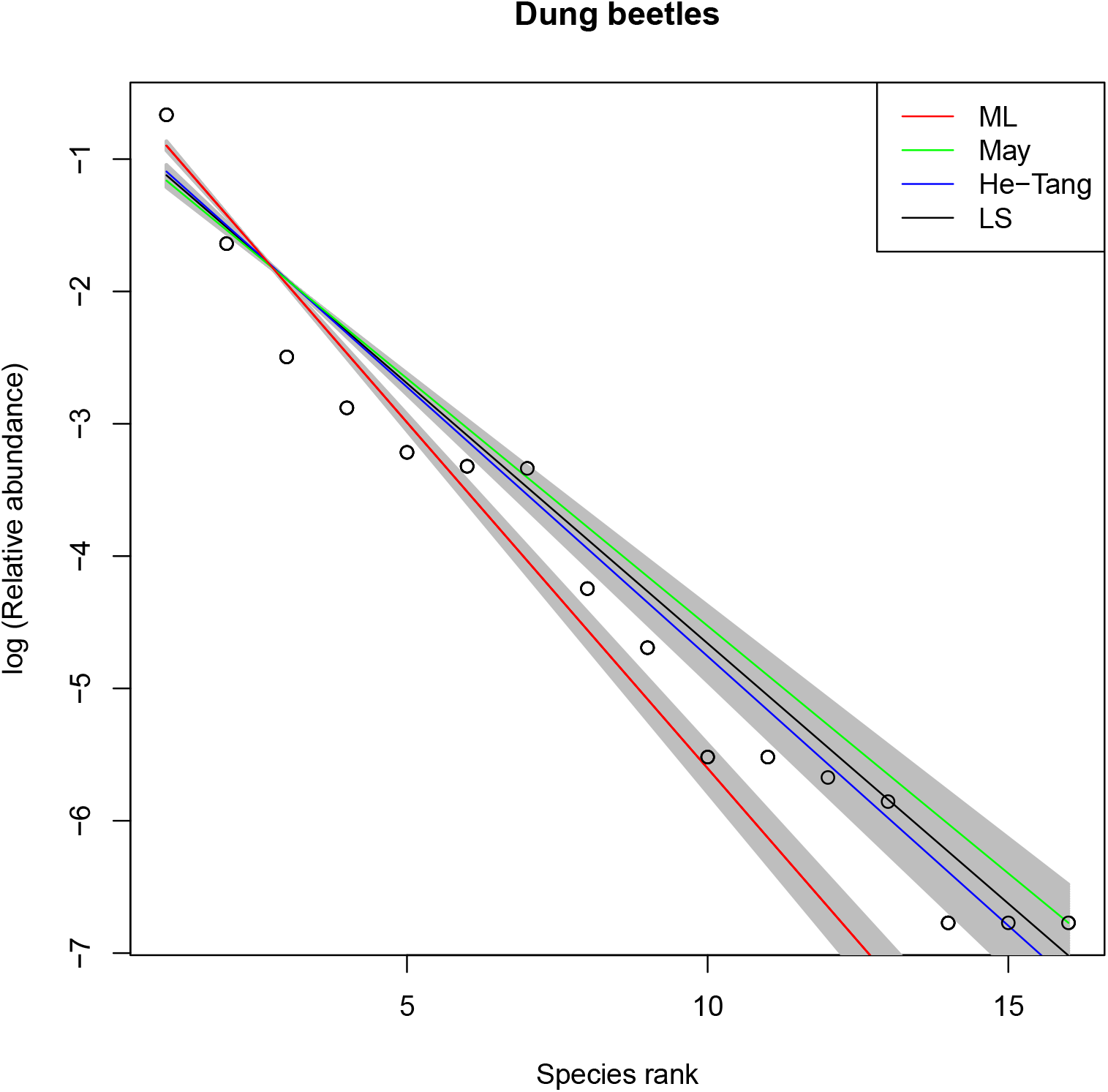
Rank-abundance plot of Indian dung beetles. Lines represent geometric decay according to four different estimators. Grey areas indicate the confidence regions for the ML and LS estimators.

**Table 3:** Estimates of the preemption parameter for the Indian dung beetles according to different methods.

| Estimator | $\hat{k}$ | se | 95% CI |
| --- | --- | --- | --- |
| May | 0.312 | - | - |
| HT | 0.334 | - | - |
| LS | 0.325 | - | (0.296, 0.353) |
| ML | 0.407 | 0.0076 | (0.392, 0.422) |

#### 4.3 Preemption *t* test with Costa Rican dung beetles

Mehrabi et al. (2014) performed a comparitive biodiversity study, where the counts of dung beetles, an important indicator taxon, were registered along eight transects under two conditions, micro-habitat standardized placement (treatment) and random placement (control) of baited traps. We use the transect level counts obtained by summing over traps sampled under the same condition. It is of interest two compare estimates of diversity parameters under the two conditions. Figure 5 shows the rank-abundance plots for the eight transects where the preemption parameter has been estimated for both conditions. Table 4 shows the ML estimates of the preemption parameter, and the results of the preemption *t* test described in Section 2. Figure 5 shows overlapping confidence intervals for transect pairs C-D, I-J, K-L and M-N. The preemption *t* test results in Table 4 show non-significant differences and overlapping confidence intervals for the first three of these, and a borderline p-value for transect M-N. All other transect pairs have very small p-values, indicating signigicant differences in the preemption parameter for the two conditions. For these transects, the ML estimator gives a faster decay for the control transects. This corroborates the finding of Mehrabi et al. (2014) that the micro-habitat standardized transects were more diverse.

**Figure 5:**
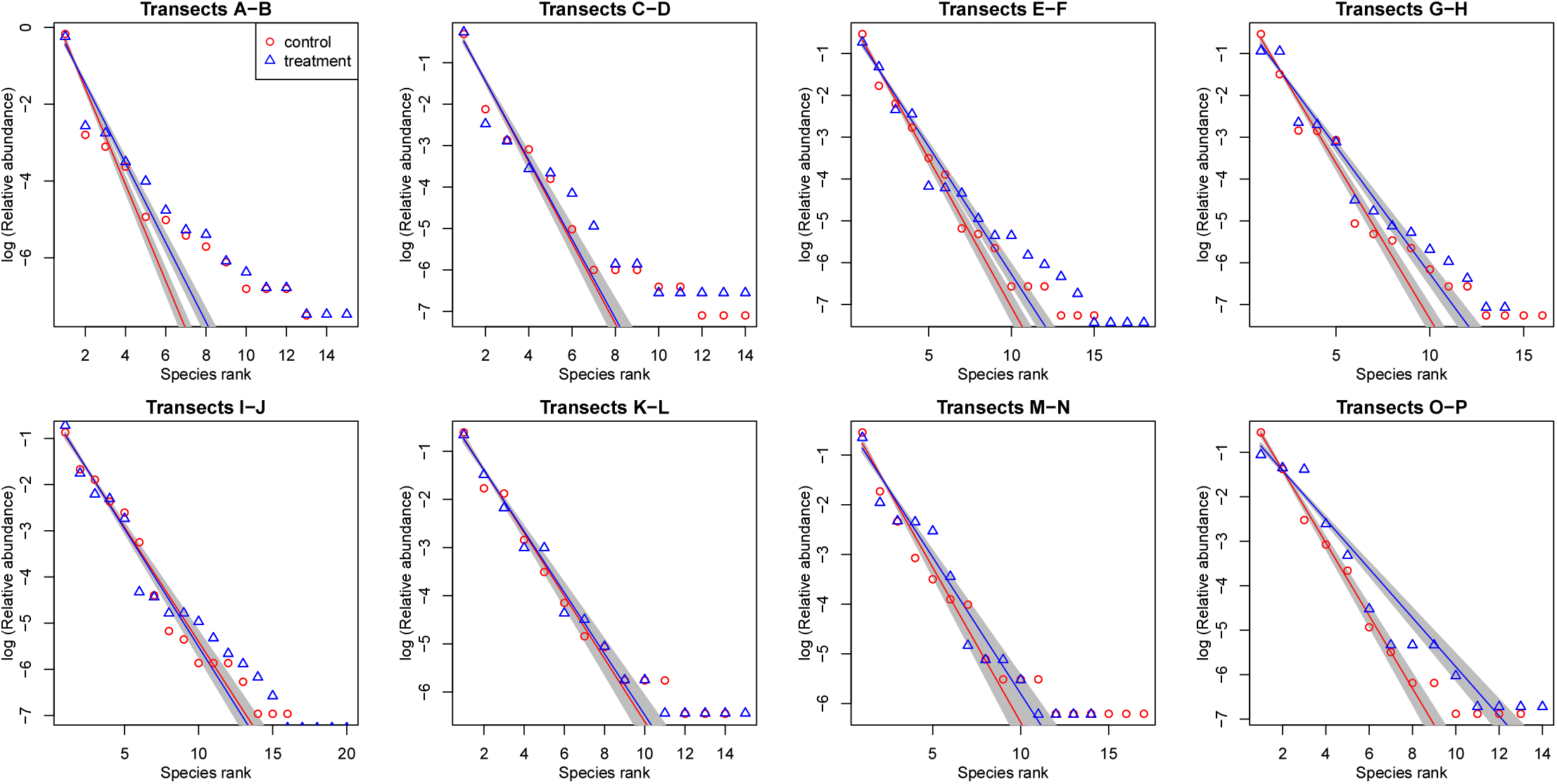
Rank-abundance plots of Costa Rican dung beetles. Lines represent geometric decay for the ML estimators for two transects, one in blue (treatment) and one in red (control). Grey areas indicate the confidence regions for the ML estimators.

**Table 4:** ML estimates 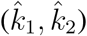 of preemption parameter *k* for eight pairs of transects under two conditions, *T*-statistic and p-value of a preemption *t*-test.

| Transects | $\hat{k}_1$ | $CI(k_1)$ | $\hat{k}_2$ | $CI(k_2)$ | $T$ | $p$-value |
| --- | --- | --- | --- | --- | --- | --- |
| A-B | 0.715 | (0.697 , 0.732) | 0.644 | (0.626 , 0.662) | 5.480 | 0.000 |
| C-D | 0.623 | (0.601 , 0.644) | 0.615 | (0.587 , 0.643) | 0.414 | 0.679 |
| E-F | 0.508 | (0.490 , 0.527) | 0.457 | (0.441 , 0.473) | 4.113 | 0.000 |
| G-H | 0.525 | (0.506 , 0.544) | 0.456 | (0.437 , 0.475) | 5.012 | 0.000 |
| I-J | 0.391 | (0.373 , 0.409) | 0.400 | (0.384 , 0.416) | -0.704 | 0.481 |
| K-L | 0.480 | (0.453 , 0.507) | 0.471 | (0.444 , 0.498) | 0.475 | 0.635 |
| M-N | 0.464 | (0.434 , 0.494) | 0.423 | (0.394 , 0.451) | 1.979 | 0.048 |
| O-P | 0.559 | (0.536 , 0.582) | 0.424 | (0.402 , 0.446) | 8.256 | 0.000 |

### 5 Conclusions and discussion

We have developed a maximum likelihood estimator for the niche preemption parameter of the geometric model. In this work, we assume the number of species *S* to be known, such that multinomial sampling with a fixed number of categories applies. In empirical studies, fixing *S* maybe reasonable if the community of interest has been exhaustively surveyed, and the number of species is known in advance. The geometric model has been found adequate for species-poor communities (Magurran, 2004) such as those in process of colonization (He and Tang, 2008). In such circumstances, the number of competing species may indeed very well be known, and fixing *S* then seems reasonable. Importantly, in such a design, zero abundances are admitted, because not all species are observed, due to the fact that some are rare, or not present in the sample by mere chance. The proposed ML estimator can deal with zeros, as the latter arise naturally under the multinomial distribution. May’s classical estimator cannot cope with zeros, as these lead to *k* = 0 or *k* = 1. He and Tang’s estimator can neither be used, because it will always produce *k* = 1 if a zero is present. Estimation by regression with log transformed relative abundances neither works for giving ln (0) = − ∞ for zero counts. Indeed, to sensibly apply all the classical estimators, zeros must first be discarded, and *S* reduced correspondingly. For the ML estimator, zeros are unproblematic. The approximate form (Eq. (9)) gives the same estimate with and without zeros, and the exact form (Eq. (8)) typically shows only minor variation due to different *S* under removal or inclusion of zeros. In future work, the maximum likelihood approach presented here could be extended to the double geometric model from Alroy (2015).

In comparative biodiversity studies, multiple samples of similar communities are often obtained. In order to sensibly do so, the same set of species is typically determined for all samples. It easily occurs that some of the rarer species are absent in some of the sampled sites. In order to apply the classical estimators, the zeros must be discarded, and consequently *S* starts to vary over the samples. To keep the same *S* constant, one can subset the analysis to those species that appear in all samples, but this obviously entails a loss of information. The ML estimator is based on the multinomial distribution and admits zeros, neatly avoiding these problems.

The least-squares regression estimator (He and Tang, 2008; Caruso and Migliorini, 2006; Fattorini, 2005) is popular, and intuitively appealing, but it suffers from certain inconsistencies. Importantly, the geometric model has, for given *N* and *S*, only one parameter, the preemption parameter *k*. However, linear regression estimates two parameters, slope *β*_1_ and intercept *β*_0_. Eq. (5) estimates *k* from the slope, but that may be considered arbitary. Because *β*_0_ also depends on *k* (see Eq. (4)), an alternative estimator for *k*, which will typically give a different point estimate, can be obtained from the intercept. Drawing a standard least-squares regression line with intercept *b*_0_ and slope *b*_1_ in the rank-abundance plot will often give a line that visually fits the data well, but it amounts to *overfitting* because the model of interest has in fact only one parameter. May’s, He-Tang’s and the proposed ML estimator are more coherent for estimating a single parameter.

We also note that the line fitted by May’s methods always passes through (*S*, ln (*n*_(*S*)_/*N*)), thereby always artifically fitting the most rare species without error. Consequently, May’s method capitalizes on the rare species. The probabilities of occurrence of the rare species are poorly estimated, because only a few individuals of them have been observed. Describing the geometric decay with an ordinary least squares regression (Eq. (4)) gives the *same weight* to highly abundance species whose proportion is determined with small relative error as to rare species whose proportion is determined with high relative error, and that looks at least questionable. The proposed ML estimator capitalizes on the abundant species and is less affected by the rare ones. The ML estimator therefore focusses on those measurements that have less relative error, which is a desirable property. If one singleton of an additional species is found, the numerator of Eq. (9) increases by 1 and the denominator by *S* + 1, which will in general hardly affect the ML estimate of *k*, showing clearly its *robustness* to the inclusion or deletion of some rare species.

There are many ecological studies in which the preemption parameter of the geometric series is estimated and reported, but a quantification of the uncertainty in the estimate is almost never given. The derivation of the ML estimator and its distribution in this article enable, by means of confidence intervals, the expression of the uncertainty in the estimation of the preemption parameter, and the comparison of such estimates by means of the preeemption *t* test.

### 6 Software

An R package (R Development Core Team, 2004) named `MLpreemption` has been written providing functions for estimation of the preemption parameter by maximum likelihood and other methods. The package also includes the preemption *t* test and the datasets analysed in this paper, and is available on CRAN and on the author’s homepage.

## 7 Acknowledgements

This work was partially supported by grants RTI2018-095518-B-C22 of the Spanish Ministry of Science, Innovation and Universities and the European Regional Development Fund, and by grant R01 GM075091 from the United States National Institutes of Health.

